# The KRAS^G12C^ Inhibitor Divarasib Stabilizes RBM39 and Antagonizes Aryl-Sulfonamide Degraders

**DOI:** 10.64898/2026.08.16.745134

**Authors:** Ssu-Yu Chen, You Zou, Jianli Wu, Gibeom Nam, Hyejin Lee, Yubin Chen, Chiara Federico, Yasaman Setayeshpour, Chao-Chieh Lin, Shao-Chin Wu, John H. Strickler, Jiyong Hong, Michael C. Fitzgerald, Jen-Tsan Chi

**Affiliations:** Department of Molecular Genetics and Microbiology, Duke University School of Medicine, Durham, NC 27710, USA; Institute of Biochemistry, National Defense Medical University, Taipei 114, Taiwan; Department of Pharmacology and Cancer Biology, Duke University School of Medicine, Durham, NC 27710, USA; Department of Chemistry, Duke University, Durham, NC 27708, USA; Cancer and Immunology Research Center, National Yang Ming Chiao Tung University, Taipei 112, Taiwan; Duke University Medical Center, Durham, NC, USA; Duke Center for Genomic and Computational Biology, Duke University School of Medicine, Durham, NC, USA

## Abstract

KRAS^G12C^ inhibitors have demonstrated meaningful clinical benefit in KRAS^G12C^-mutant non-small cell lung cancer (NSCLC), yet responses remain heterogeneous and treatment-associated toxicities persist for reasons that are incompletely understood. Cysteine profiling indicates that these covalent inhibitors are highly selective for mutant KRAS; however, such approaches cannot detect noncovalent engagement of additional non-RAS proteins. Here, we used a protein-folding stability profiling technique, stability of proteins from rates of oxidation (SPROX), to identify protein targets of the clinical KRAS^G12C^ inhibitor, divarasib (GDC-6036), in KRAS-mutant NSCLC lysates. SPROX revealed a focused set of candidate interactors, including the essential splicing factor RBM39, which was reproducibly stabilized at both divarasib concentrations tested. We subsequently confirmed that divarasib directly and noncovalently binds to RBM39 protein. In NSCLC cells, divarasib increased RBM39 protein abundance and antagonized RBM39 degradation induced by the aryl-sulfonamide molecular glue indisulam through a post-transcriptional mechanism. Divarasib and RBM39 degraders reciprocally antagonized each other’s cytotoxicity, and RBM39 knockdown modestly reduced divarasib-induced cell death. Mechanistically, divarasib-mediated RBM39 stabilization regulated both INSR expression and alternative splicing, altered downstream insulin receptor signaling, and contributed to divarasib-associated cytotoxicity. Consistent with these findings, RBM39 and INSR expression were positively correlated across multiple human cancer types. Collectively, these findings identify RBM39 as a previously unrecognized noncovalent target of divarasib and uncover an RBM39–INSR signaling axis that modulates cellular responses to both divarasib and RBM39 degraders.

**Significance:** Covalent KRAS^G12C^ inhibitors are widely considered highly target-selective because adduct-based chemoproteomics approaches almost exclusively identify mutant KRAS as their target, detecting few proteins beyond it. However, these methods cannot capture noncovalent interactions and can sometimes miss covalent interactors. Using stability-based proteomics, we show that the clinical KRAS^G12C^ inhibitor divarasib targets RBM39, an essential splicing factor degraded by anticancer aryl-sulfonamide molecular glues. Divarasib opposes degrader-induced RBM39 loss, attenuates degrader cytotoxicity, and preserves RBM39-dependent regulation of the insulin receptor. These findings uncover a previously unrecognized non-KRAS axis of divarasib activity that is invisible to conventional target-deconvolution methods. They also suggest potential liability associated with combining RBM39 degraders with divarasib or other KRAS^G12C^ inhibitors that engage RBM39. More broadly, our study illustrates how protein folding stability-based profiling can expand the actionable target landscape of small molecules beyond canonical covalent interactions, uncovering noncovalent off-targets that may underlie response heterogeneity and treatment-associated toxicity.

**HIGHLIGHTS:**

- Protein folding stability profiling identifies RBM39 as a noncovalent target of divarasib.
- Divarasib stabilizes RBM39 and blocks its degradation by aryl-sulfonamide glues.
- Divarasib and RBM39 degraders reciprocally antagonize each other’s cytotoxicity.
- RBM39 supports insulin receptor expression and divarasib-induced cell death.

**eTOC BLURB:** Chen et al. apply protein folding stability-based proteomics to the covalent KRAS^G12C^ inhibitor divarasib and identify the splicing factor RBM39 as a noncovalent target of divarasib. Divarasib stabilizes RBM39, antagonizes aryl-sulfonamide degraders, and preserves RBM39-dependent insulin receptor signaling, revealing a non-KRAS axis invisible to adduct-based target deconvolution.

## Introduction

The RAS family of genes (HRAS, KRAS, and NRAS) encodes small GTPases that regulate cellular proliferation, differentiation, and survival. Under physiological conditions, RAS proteins act as molecular switches that cycle between an active GTP-bound state and an inactive GDP-bound state. Point mutations in RAS genes, particularly at positions G12, G13, or Q61, result in constitutively active RAS proteins that drive growth-promoting signaling independent of extracellular stimuli (1). Among these alterations, mutant KRAS (mKRAS) represents the most prevalent oncogenic RAS lesion in pancreatic, colorectal, and lung cancers and is associated with poor therapeutic responses. Consequently, the development of strategies to target mKRAS, including efforts led by the National Cancer Institute (NCI) RAS Initiative (2), has been a longstanding priority in cancer research.

mKRAS was long regarded as “undruggable” until the development of selective covalent inhibitors targeting the KRAS^G12C^ mutant. Sotorasib (AMG 510), the first-in-class KRAS^G12C^ inhibitor, represented a major breakthrough and received accelerated FDA approval for the treatment of KRAS^G12C^-mutated NSCLC (3, 4). This landmark drug not only overcame a longstanding challenge in cancer therapeutics but also stimulated broad interest in targeting other mutant RAS proteins. Divarasib (GDC-6036) is a second-generation covalent KRAS^G12C^ inhibitor with enhanced potency and selectivity, demonstrating promising clinical activity as monotherapy and in combination regimens (5, 6). Its clinical profile has since been reinforced by the Phase III Krascendo 1 trial (NCT06497556), the first global head-to-head comparison of KRAS^G12C^ inhibitors, in which divarasib monotherapy improved both progression-free and overall survival relative to the approved first-generation inhibitors sotorasib and adagrasib in previously treated KRAS^G12C^-mutant NSCLC (7). The success of these agents has further accelerated the development of inhibitors targeting other mKRAS alleles, including G12D-selective inhibitors such as MRTX1133 (8) and pan-KRAS inhibitors such as daraxonrasib (RMC-6236) (9). Most recently, daraxonrasib (RMC-6236), an oral multi-selective RAS (ON) inhibitor developed by Revolution Medicines, demonstrated a significant overall survival benefit in the pivotal Phase III RASolute 302 trial in patients with previously treated metastatic pancreatic ductal adenocarcinoma (10). Median overall survival was nearly doubled compared with standard chemotherapy (13.2 vs. 6.7 months), providing compelling clinical proof of concept that direct inhibition of the activated RAS can improve survival in KRAS-driven cancer.

Although mKRAS inhibitors have demonstrated remarkable clinical efficacy, therapeutic responses vary considerably even among tumors harboring the same KRAS mutation (5, 11), and their use is associated with adverse effects that are not fully accounted for by on-target KRAS inhibition (5, 12). This variability in efficacy, together with the emergence of toxicities (5), suggests that the mechanisms underlying their antitumor effects are not yet fully understood and that additional determinants of drug response, potentially including engagement of proteins beyond mKRAS, remain to be identified. Several chemoproteomic studies have reported high selectivity for these covalent inhibitors, with cysteine profiling revealing target engagement primarily at mKRAS and only a limited number of additional proteins (3, 13). However, because adduct-centric approaches detect only covalent engagement, it remains largely unexplored whether these inhibitors also bind proteins noncovalently.

Detecting such noncovalent engagement requires approaches that do not depend on covalent adduct formation. Stability-based proteomic methods provide one such strategy, detecting ligand engagement without chemical derivatization of the ligand. These approaches include thermal proteome profiling (TPP) (14), proteolysis-based strategies such as drug affinity responsive target stability (DARTS) (15), limited proteolysis (16), pulse proteolysis (17), and stability of proteins from the rates of oxidation (SPROX) (18). Because these methods measure ligand-induced changes in protein folding stability arising from either direct binding or allosteric effects, they can, in principle, detect noncovalent interactions that are inaccessible to conventional activity-based protein profiling or covalent pull-down approaches. Furthermore, “one-pot” implementations increase throughput and facilitate the identification of high-confidence interactors (19, 20). Recent proof-of-principle work from our groups has shown that such stability-based approaches do indeed add value to conventional activity-based profiling or covalent pull-down approaches (21).

To test whether the clinical KRAS^G12C^ inhibitor divarasib engages targets beyond its well-established covalent site in KRAS^G12C^, we applied a one-pot SPROX workflow to profile divarasib-induced changes in protein-folding stability in KRAS^G12C^-mutant H358 NSCLC cell lysates and identified RBM39 (RNA-binding motif protein 39) as a candidate interactor. Divarasib engagement of RBM39 in cells was confirmed by cellular thermal shift assay (CETSA), and direct binding to purified recombinant RBM39 was demonstrated by MicroScale Thermophoresis (MST). Because RBM39 was not detected as a covalent target in adduct-based profiling (22), these findings support a noncovalent mode of engagement and identify RBM39 as a previously unrecognized noncovalent target of divarasib. We further found that divarasib increased RBM39 abundance, antagonized RBM39 degradation induced by aryl-sulfonamide molecular glues, and attenuated degrader-induced cytotoxicity. Furthermore, divarasib-mediated RBM39 stabilization altered INSR expression and alternative splicing, linking the RBM39–INSR axis to insulin receptor signaling and to the cytotoxic response evoked by these compounds. Collectively, these findings establish RBM39 as a previously unrecognized protein target of divarasib and uncover an RBM39–INSR signaling axis that modulates cellular responses to both divarasib and aryl-sulfonamide degraders in KRAS-mutant NSCLC.

## Results

### SPROX identifies candidate divarasib targets in NSCLC

To profile divarasib-induced changes in protein stability, we applied a one-pot SPROX workflow to H358 NSCLC cell lysates where we quantified methionine oxidation across a denaturant gradient using a multiplexed bottom-up proteomics approach (18). Briefly, lysates from biological replicates were treated with vehicle (DMSO), 1 μM divarasib, or 100 μM divarasib. The 1 μM concentration was selected based on the clinically relevant peak plasma concentration reported previously (5). In SPROX, the magnitude of protein stability changes is proportional to the free ligand concentration (18). Therefore, an additional high-concentration condition (100 μM) was included to enhance the detection of stability changes in protein–ligand complexes with weaker binding affinities. Samples were subsequently exposed to a urea denaturation gradient, followed by H₂O₂-mediated methionine oxidation (23, 24). Reactions were quenched with TCEP, pooled into a single one-pot sample for each biological replicate, and analyzed by quantitative bottom-up proteomics using tandem mass tag (TMT) labeling. Proteins exhibiting divarasib-induced stability changes were identified as SPROX hits based on Welch’s *t*-test (*P* < 0.05) and an absolute log_2_ fold-change threshold of 0.3 (Figure 1A).

**Figure 1.**
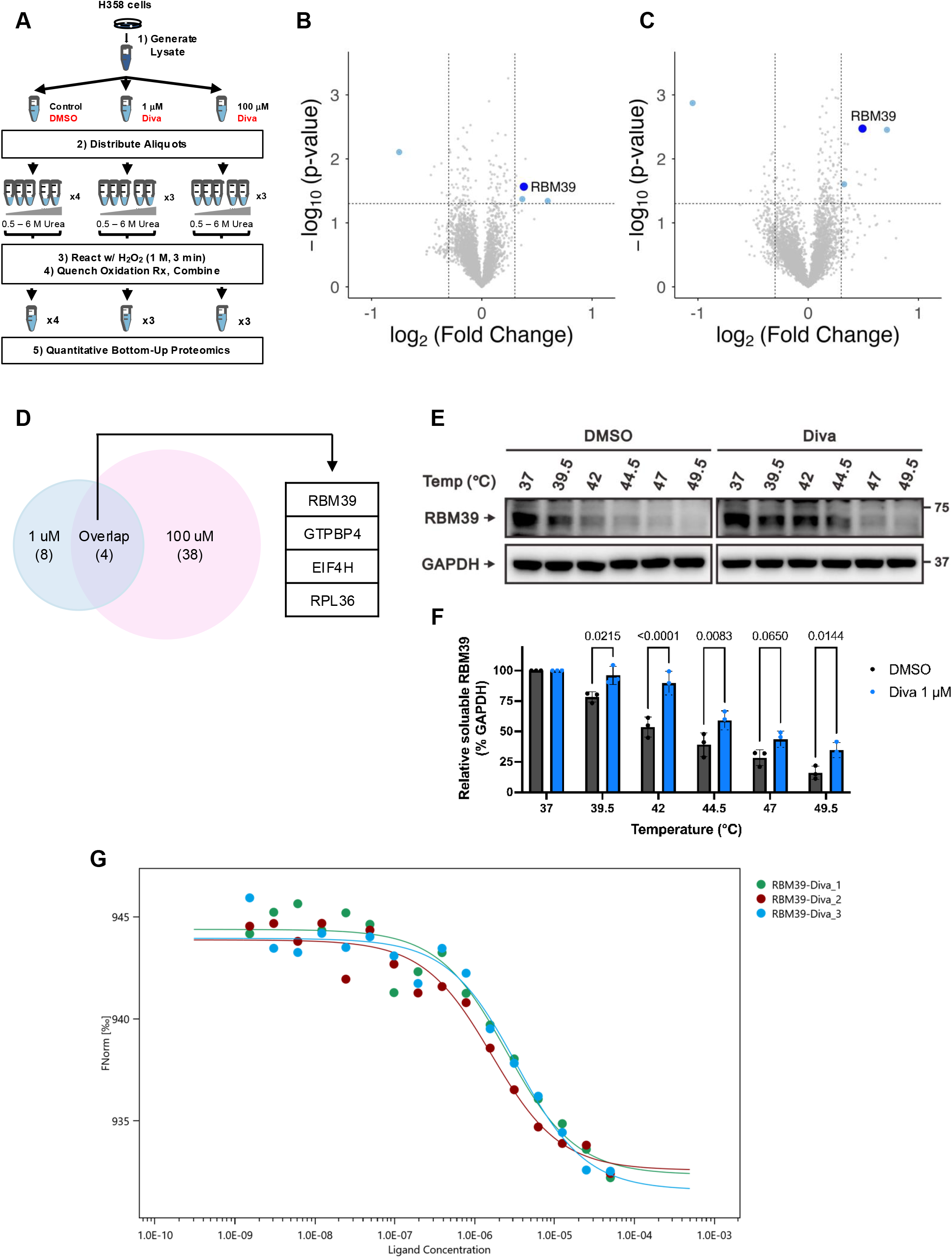
One-pot SPROX analysis reveals divarasib-induced protein folding stability changes in H358 cell lysates. (A) Schematic illustration of the one-pot SPROX workflow. H358 cell lysates were treated with DMSO, 1 μM divarasib (Diva), or 100 μM Diva, followed by incubation in a urea denaturation gradient and oxidation with H₂O₂. Reactions were quenched with TCEP and combined into a single “one-pot” sample for subsequent proteomic analysis. SPROX was performed with four biological replicates for the DMSO condition and three for each divarasib condition. (B, C) Volcano plots of one-pot SPROX results comparing (B) 1 μM Diva and (C) 100 μM Diva with DMSO-treated controls. (D) Venn diagram showing the overlap of significant hits identified in both the 1 μM (12 hits) and 100 μM (42 hits) Diva-treated samples. Four proteins, RBM39, GTPBP4, EIF4H, and RPL36, were shared between both conditions. Significant hits were defined by Welch’s t-test (p < 0.05) and |log_2_FC| > 0.3. (E, F) Cellular thermal shift assay (CETSA) performed in H358 cells treated with DMSO or 1 μM Diva for 1 h and subjected to a temperature gradient ranging from 37 °C to 49.5 °C. (E) Representative immunoblots showing soluble RBM39 and GAPDH levels. (F) Quantification of soluble RBM39 normalized to GAPDH at each temperature. Data are presented as mean ± SD from three independent biological replicates. (G) MicroScale Thermophoresis (MST) analysis of divarasib binding to recombinant RBM39. Divarasib was titrated against 50 nM RED-tris-NTA-labeled His-RBM39 in PBS-T. Points represent individual measurements from three independent experiments (green, red, and blue), and lines represent fits to a 1:1 binding model with the target concentration fixed at 50 nM. Kd = 2.5 ± 0.9 µM (mean ± SD, n = 3). Statistical significance in (F) was determined by ordinary two-way ANOVA followed by Šídák’s multiple-comparisons test.

Using the above statistical thresholds, we identified a focused set of candidate proteins exhibiting divarasib-dependent stability shifts (Supplementary Table 1). At 1 μM divarasib, SPROX identified 12 methionine-containing peptides mapping to 12 proteins that exhibited significant stability changes (Figure 1B). At 100 μM divarasib, 43 methionine-containing peptides representing 42 proteins met the hit criteria (Figure 1C), with 4 proteins shared between the two concentrations (Figure 1D). Among the overlapping hits was RBM39 (RNA-binding motif protein 39), an RNA splicing factor and established therapeutic target implicated in multiple cancers (25–28). Another shared hit, GTPBP4 (guanosine triphosphate-binding protein 4), is frequently upregulated in tumors and has been linked to cell cycle progression and MAPK signaling (29–31). EIF4H (eukaryotic translation initiation factor 4H) participates in translation initiation (32, 33), whereas RPL36 (ribosomal protein L36) is a component of the 60S ribosomal subunit involved in ribosome assembly and protein synthesis. Given its established role in RNA splicing and therapeutic relevance, RBM39 was prioritized for further validation and mechanistic studies.

Proteome-wide cysteine-reactivity profiling showed that divarasib competes only the KRAS^G12C^ cysteine among more than 11,000 quantified cysteines, providing no evidence of covalent engagement at RBM39 (22). Consistent with this, a second independent reactive-cysteine profiling study that assessed divarasib against more than 22,000 cysteines in the same KRAS^G12C^-mutant H358 background likewise identified KRAS^G12C^ as the dominant covalent target (34). Because the second study (34) deposited its raw cysteine-level data, we examined the RBM39 peptides directly. Of the five cysteines in RBM39 (C157, C303, C425, C454, and C468), two were detected, C157 and C303, and neither was covalently engaged by divarasib (Supplementary Figure 1A). Notably, the SPROX-responsive peptide reporting on RBM39 stability (M184; residues 184– 191) localizes to the first RNA-recognition motif (RRM1), which is close in primary structure to one of the RBM39 cysteines covered in that dataset (C157) but remote from the three RBM39 cysteines not covered in that dataset (C425, C454, and C468). Together, these orthogonal cysteine-profiling datasets argue against covalent modification of RBM39 within the RRM1 region whose folding stability is altered by divarasib and support a noncovalent mode of engagement.

To further assess target engagement in cells, we performed CETSA (35). H358 cells were treated with vehicle (DMSO) or 1 μM divarasib and subjected to a temperature gradient. Heat-induced protein aggregation decreases the soluble protein fraction, whereas ligand binding can stabilize target proteins and increase their thermal resistance. Soluble RBM39 levels were subsequently measured by immunoblotting. Consistent with the SPROX results, divarasib treatment increased RBM39 thermal stability at most temperatures tested between 39.5 °C and 49.5 °C, providing further evidence that divarasib engages RBM39 in cells (Figure 1E–F). We next asked whether divarasib directly binds RBM39 using MST. Titration of divarasib (1.53 nM– 50 μM) against 50 nM RED-tris-NTA-labeled recombinant His-tagged RBM39 (Thr146-Val326) produced a saturable, concentration-dependent change in normalized thermophoresis (mean response amplitude 11.9 ± 0.5 FNorm units; mean signal-to-noise 14.4). Across three independent experiments, fitting the binding curves to a 1:1 binding model yielded an apparent dissociation constant of 2.5 ± 0.9 μM (mean ± SD), demonstrating direct, noncovalent binding of divarasib to RBM39 protein (Figure 1G). Together, the cysteine-reactivity profiling, SPROX, CETSA, and MST results all support RBM39 as a noncovalent target of divarasib.

To investigate the molecular basis of divarasib interaction with RBM39, we performed computational docking analysis, focusing on the RRM1 domain, which contains the SPROX-responsive peptide (MISDRNSR; residues 184–191). To date, the full structure of RBM39 has not been determined. However, NMR spectroscopy-based structures of RRM1 and RRM2 domains complexed with RNA have been reported (36). The SPROX-responsive peptide corresponds to the β2–β3 loop of the RRM1 domain, which is involved in RNA hairpin recognition and is essential for the splicing function of RBM39 (Supplementary Figure 1B) (36). We therefore hypothesized that divarasib binds within the groove formed by the β2–β3 loop and the C-terminal region of the RRM1 domain and preformed docking without the RNA structure. Docking predicted a pose in which divarasib occupies the groove, with hydrogen bonds to Ser190, Arg192, and Gln226, suggesting potential binding of divarasib to the RRM1 domain of RBM39 (Supplementary Figure 1C and D). Collectively, these results suggest that RBM39 may represent a previously unrecognized noncovalent target of divarasib.

### Divarasib increases RBM39 abundance and antagonizes RBM39 degraders in NSCLC

Having identified RBM39 as a divarasib interactor, we next investigated how divarasib affects RBM39 expression and function. In H358 cells, divarasib treatment resulted in a marked band shift of KRAS protein due to covalent binding to KRAS^G12C^ mutations (Figure 2A). We also found that basal RBM39 protein levels were slightly but significantly increased after 72 hours of divarasib treatment (Figure 2A and C). Conversely, RBM39 mRNA levels remained unchanged following divarasib treatment (Figure 2E), suggesting that the increase in RBM39 protein abundance is mediated through post-transcriptional mechanisms, potentially through enhanced protein stability. Consistent with these findings, divarasib-induced upregulation of RBM39 was also observed in another KRAS^G12C^-mutant NSCLC cell line, H2122, at both concentrations tested (Supplementary Figure 2A–B).

**Figure 2.**
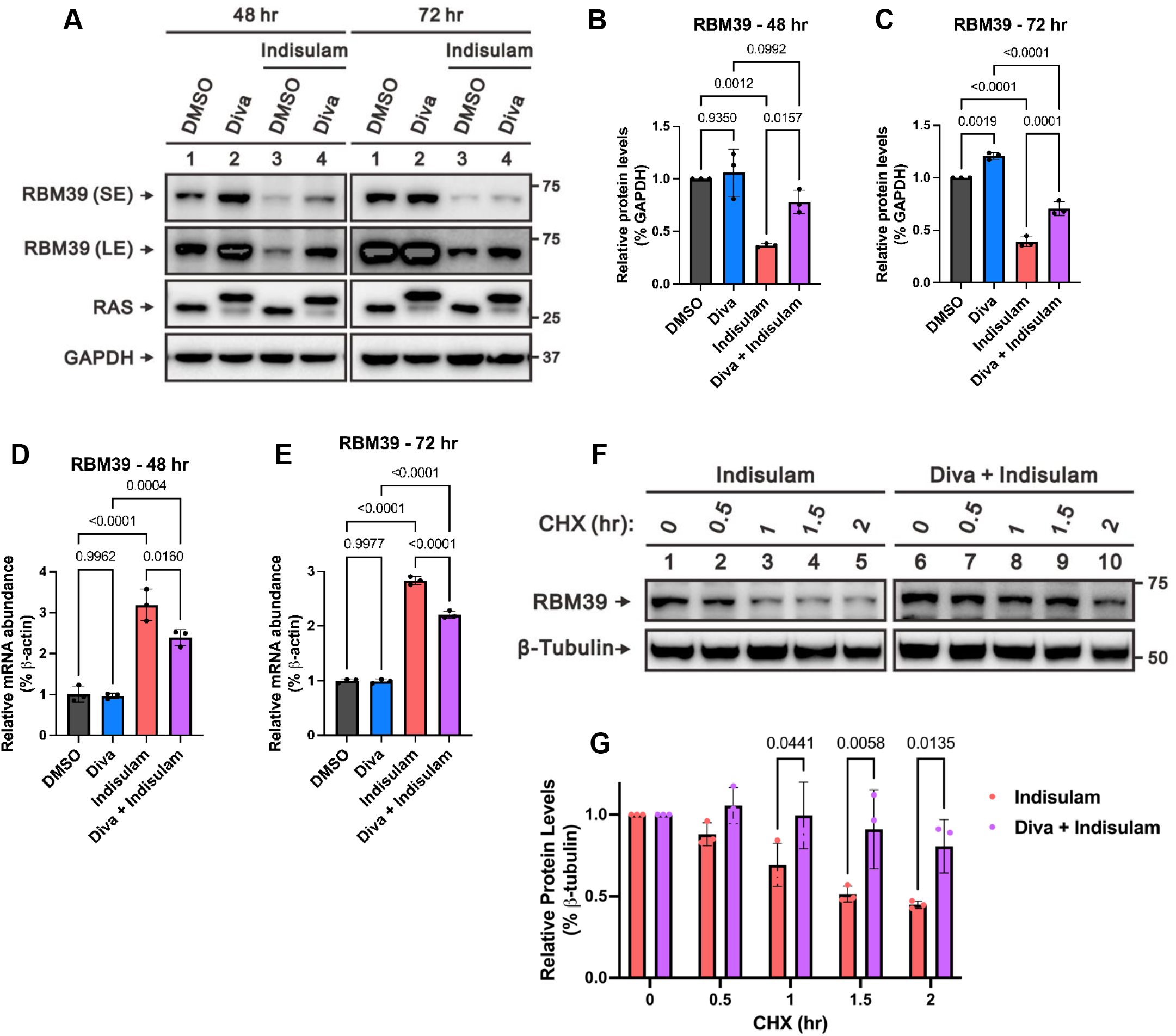
Divarasib increases RBM39 abundance and antagonizes RBM39 degraders in H358 cells. (A-C) RBM39 protein levels in H358 cells treated with DMSO, divarasib (Diva, 4 nM), indisulam (5 μM), or the combination of Diva and indisulam for 48 or 72 h. (A) Representative immunoblots of RBM39 at short (SE) and long (LE) exposure, RAS, and GAPDH. (B, C) Quantification of RBM39 protein levels normalized to GAPDH at (B) 48 h or (C) 72 h. (D, E) RBM39 mRNA expression levels in H358 cells treated with DMSO, Diva (4 nM), indisulam (5 μM), or the combination of Diva and indisulam for (D) 48 h or (E) 72 h, as determined by RT-qPCR. mRNA levels were normalized to β-actin. (F–G) RBM39 protein stability in H358 cells treated with indisulam (5 μM) or the combination of Diva (4 nM) and indisulam for 12 h, followed by a cycloheximide chase for 0-2 h. (F) Representative immunoblots of RBM39 and β-Tubulin. (G) Quantification of RBM39 protein levels normalized to β-Tubulin. Data are presented as mean ± SD from three independent biological replicates. Statistical significance in (B-E) was determined by ordinary one-way ANOVA followed by Tukey’s multiple-comparisons test. Statistical significance in (G) was determined by ordinary two-way ANOVA followed by Šídák’s multiple-comparisons test.

Aryl-sulfonamide molecular glues, such as indisulam, are a class of RBM39 degraders that induce RBM39 degradation by recruiting RBM39 to the DCAF15 E3 ubiquitin ligase (37, 38). We first confirmed that indisulam substantially reduced RBM39 protein levels in H358 cells (Figure 2A–C) and induced a compensatory increase in RBM39 mRNA levels (Figure 2D–E), validating its on-target activity. We then sought to determine how divarasib, which was found to interact with RBM39, would affect RBM39 protein levels under degrader treatment. Surprisingly, co-treatment with divarasib significantly restored RBM39 protein abundance in the presence of indisulam at both 48 and 72 hours (Figure 2A–C). Importantly, divarasib significantly reduced the indisulam-induced increase in RBM39 mRNA at both 48 and 72 hours (Figure 2D–E) rather than further elevating it, indicating that the rescue of RBM39 protein occurs through a post-transcriptional mechanism and is accompanied by relief of the compensatory transcriptional response. Consistent with this interpretation, cycloheximide chase experiments demonstrated that divarasib significantly increased RBM39 protein stability compared with indisulam treatment alone (Figure 2F–G). Because divarasib directly binds RBM39, one possible explanation is that this interaction may interfere with degrader-induced RBM39 turnover. Whether this reflects competition at the DCAF15 interface, an allosteric effect transmitted from the RRM1 region, or an indirect mechanism remains to be determined. Taken together, these results indicate that divarasib stabilizes RBM39 at the protein level, linking its direct binding and cellular target engagement (Figure 1) to a functional consequence—protection of RBM39 from degrader-induced loss.

### Divarasib antagonizes the cytotoxic effects of RBM39 degraders in NSCLC

Divarasib is a covalent KRAS^G12C^ inhibitor that induces significant cell death in NSCLC cells harboring KRAS^G12C^ mutations (5). Indisulam has also demonstrated therapeutic potential across several cancer types (39–42) and has been evaluated in multiple clinical trials, in which it was shown to be safe and well tolerated (43–51). Since we found that divarasib and indisulam exert opposite effects on RBM39 protein levels, we next investigated how this regulation influences the therapeutic efficacy of these two agents in NSCLC. Consistent with our previous findings, divarasib attenuated the cytotoxic activity of indisulam in a dose-dependent manner in H358 cells (Figure 3A). To further validate this antagonistic interaction, we included two additional RBM39 degraders, E7820 and dCeMM1 (52). Similarly, divarasib exhibited strong protective effects against cell death induced by both compounds (Figure 3B–C). Reciprocally, all three RBM39 degraders significantly rescued cells from divarasib-induced cell death in a dose-dependent manner (Figure 3D–F). These reciprocal antagonistic effects suggest that opposite regulation of RBM39 protein abundance contributes to the distinct cytotoxic activities of divarasib and RBM39 degraders.

**Figure 3.**
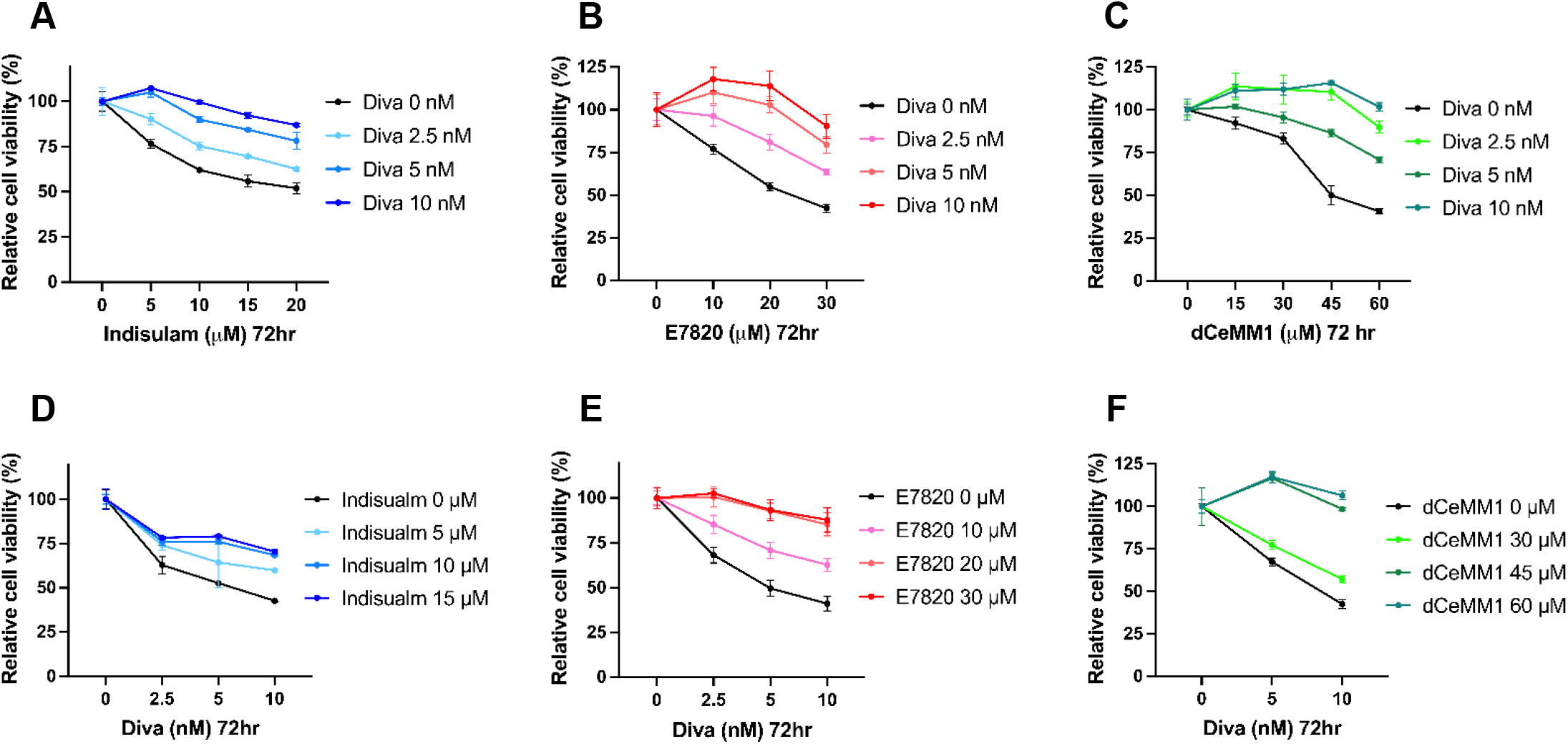
Divarasib and RBM39 degraders mutually antagonize cytotoxicity in H358 cells. (A-C) Cell viability of H358 cells was determined by CellTiter-Glo assay following treatment with increasing concentrations of the RBM39 degraders (A) indisulam, (B) E7820, or (C) dCeMM1 in the presence of the indicated fixed concentrations of divarasib (Diva) for 72 h. (D-F) Cell viability of H358 cells was determined by CellTiter-Glo assay following treatment with increasing concentrations of Diva in the presence of the indicated fixed concentrations of (D) indisulam, (E) E7820, or (F) dCeMM1 for 72 h. Data are presented as mean ± SD from three independent biological replicates. Statistical significance was determined by ordinary two-way ANOVA followed by Tukey’s multiple-comparisons test for panels (A-F).

### Divarasib-induced RBM39 stabilization implicates INSR signaling as a downstream pathway in NSCLC

RBM39 is a serine/arginine-rich RNA-binding protein that functions as a transcriptional coactivator and a key regulator of pre-mRNA splicing (25, 53). To further investigate how divarasib affects RBM39 function, we sought to identify downstream gene expression changes associated with divarasib-mediated RBM39 stabilization. To this end, we performed RNA-sequencing on H358 cells treated with divarasib or vehicle control (Figure 4A). Given that divarasib increased RBM39 protein abundance, we intersected genes significantly upregulated by divarasib treatment (log_2_ fold change > 1.5; 1,064 genes) with previously reported RBM39-regulated splicing targets (359 genes) (54). This analysis identified five candidate genes whose expression or splicing may be influenced by divarasib-induced RBM39 stabilization (Figure 4B). To further prioritize these candidates, we interrogated The Cancer Genome Atlas (TCGA) datasets and identified genes whose expression positively correlated with RBM39 in both lung adenocarcinoma (LUAD) and lung squamous cell carcinoma (LUSC). Among these candidates, INSR and CACNB3 exhibited positive correlations with RBM39 expression (Figure 4C-D and Supplementary Figure 3). Moreover, RBM39 and INSR expression showed positive correlations across most TCGA tumor types (Figure 4E), suggesting that the association between these genes extends beyond lung cancer. Consistent with these findings, Gene Set Enrichment Analysis (GSEA) of divarasib-treated cells revealed enrichment of insulin receptor recycling pathways (Figure 4F), further supporting a role for insulin receptor signaling in the cellular response to divarasib. Based on the strength of the association and the pathway enrichment analysis, we selected the insulin receptor gene (INSR) for further mechanistic investigation.

**Figure 4.**
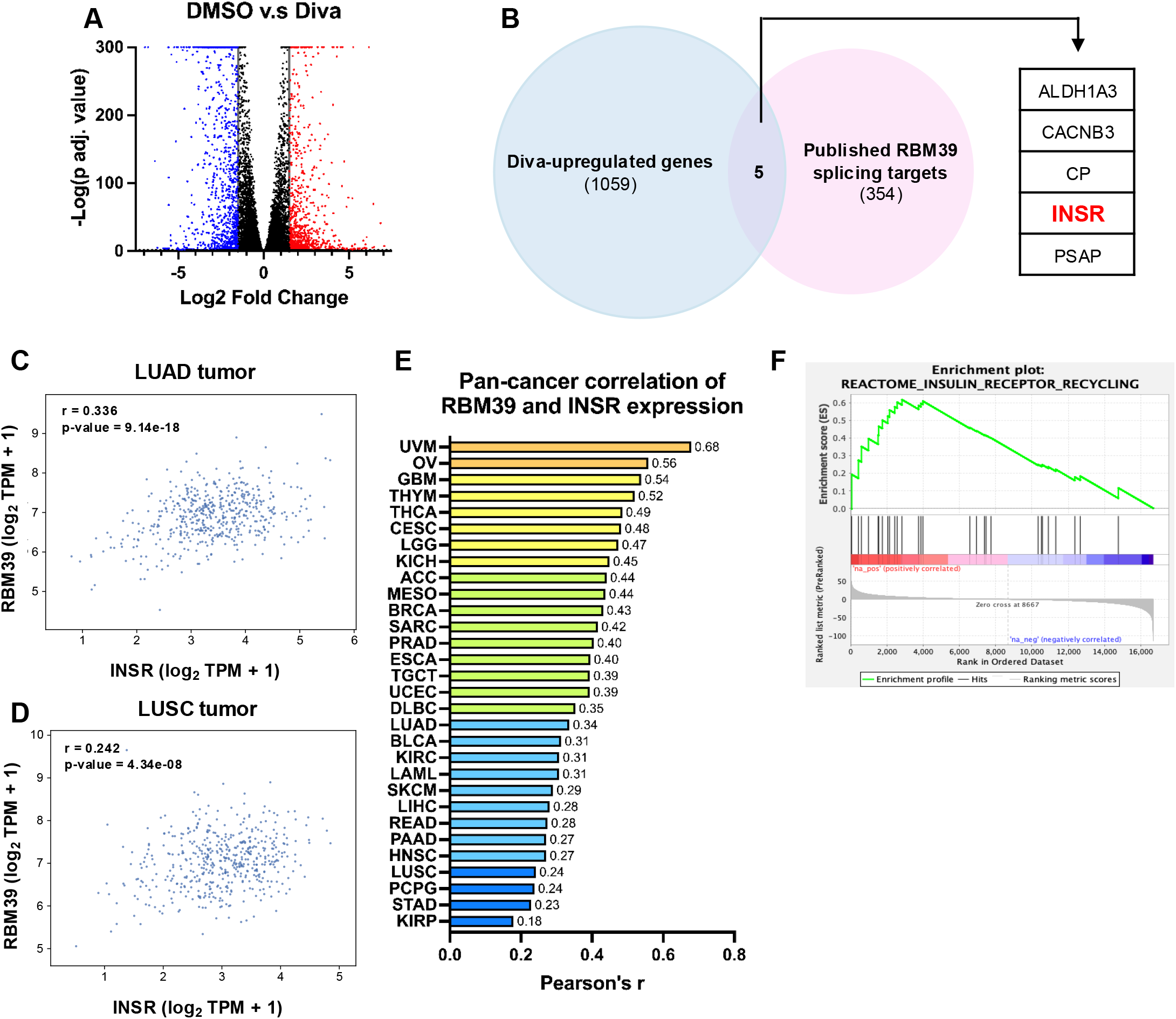
Divarasib-induced RBM39 stabilization implicates INSR signaling in H358 cells. (A) Volcano plot of RNA-seq differential expression analysis comparing DMSO- and divarasib (Diva)-treated H358 cells. (B) Overlap between Diva-upregulated genes (1,064) and previously reported RBM39-regulated splicing targets (359) (54), identifying five shared candidate genes (ALDH1A3, CACNB3, CP, INSR, and PSAP). (C–E) Correlation between RBM39 and INSR expression across TCGA cancer datasets. Scatter plots show RBM39 versus INSR transcript levels (log2 TPM + 1) in (C) lung adenocarcinoma (LUAD) and (D) lung squamous cell carcinoma (LUSC), while (E) summarizes Pearson’s correlation coefficients across the 30 cancer types exhibiting significant positive correlations. Data shown in (C–E) are based on data generated by the TCGA Research Network (https://www.cancer.gov/tcga). (F) Gene Set Enrichment Analysis (GSEA) of Diva-treated H358 cells showing enrichment of the insulin receptor recycling pathway.

### Divarasib-mediated RBM39 stabilization modulates INSR expression and alternative splicing in NSCLC

The insulin receptor is a receptor tyrosine kinase composed of two extracellular ligand-binding α-subunits and two transmembrane β-subunits containing the intracellular kinase domain. Alternative splicing of exon 11 of the INSR gene gives rise to two isoforms, INSR-A and INSR-B. INSR-A lacks exon 11 and preferentially binds insulin-like growth factor 2 (IGF-2), thereby promoting mitogenic signaling, whereas INSR-B includes exon 11 and primarily mediates the metabolic actions of insulin. A shift toward expression of the INSR-A isoform has been reported in multiple cancer types and is thought to facilitate tumor growth and survival (55–57). Given the dual roles of RBM39 as a transcriptional coactivator and a regulator of pre-mRNA splicing, we next investigated whether divarasib-mediated stabilization of RBM39 affects the expression and alternative splicing of the insulin receptor. We found that divarasib treatment significantly increased total INSR mRNA levels, whereas indisulam reduced INSR expression after 48 and 72 hours of treatment. Importantly, co-treatment with divarasib reversed the decrease in INSR expression induced by indisulam, mirroring changes in RBM39 protein abundance and suggesting that RBM39 regulates INSR expression (Figure 5A-B). Consistent with these findings, insulin receptor β protein levels exhibited similar changes (Figure 5C-E). Additionally, siRNA-mediated depletion of RBM39 led to markedly reduced levels of insulin receptor β protein, further supporting a functional association between RBM39 and INSR regulation (Figure 6A). Collectively, these findings support a role for RBM39 in regulating INSR expression.

**Figure 5.**
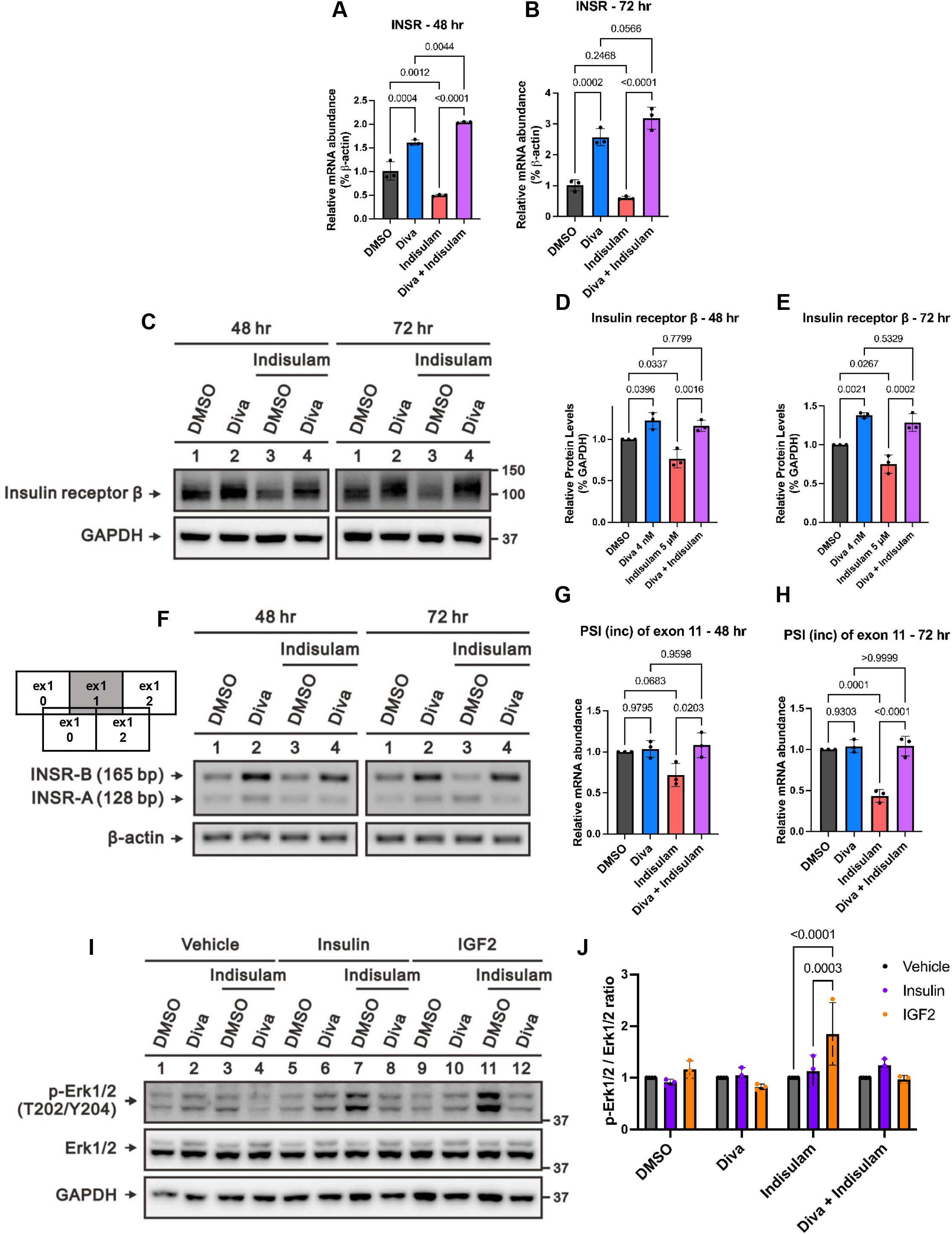
Divarasib-mediated RBM39 stabilization modulates INSR expression and alternative splicing in H358 cells. (A, B) INSR mRNA levels in H358 cells treated with DMSO, divarasib (Diva, 4 nM), indisulam (5 μM), or the combination for (A) 48 h or (B) 72 h, as determined by RT-qPCR and normalized to β-actin. (C–E) Insulin receptor β protein levels under the same treatment conditions. (C) Representative immunoblots of insulin receptor β and GAPDH at 48 and 72 h. (D, E) Quantification of insulin receptor β protein levels at (D) 48 h and (E) 72 h, normalized to GAPDH. (F–H) INSR exon 11 alternative splicing in H358 cells under the same treatment conditions. (F) Schematic representation and representative RT-PCR analysis detecting INSR-B (exon 11 included, 165 bp) and INSR-A (exon 11 excluded, 128 bp) isoforms, with β-actin as loading control. (G, H) Percent spliced in (PSI) values for exon 11 at (G) 48 h and (H) 72 h. (I, J) ERK1/2 phosphorylation at Thr202/Tyr204 in H358 cells treated with DMSO, Diva (4 nM), indisulam (5 μM), or the combination for 48 h, followed by serum starvation for 6 h and stimulation with insulin (100 nM) or IGF-2 (100 ng/mL) for 30 min. (I) Representative immunoblots of phospho-ERK1/2, total ERK1/2, and GAPDH. (J) Quantification of phospho-ERK1/2 levels normalized to total ERK1/2. Data are presented as mean ± SD from three independent biological replicates. Statistical significance in (A, B, D, E, G, and H) was determined by ordinary one-way ANOVA followed by Tukey’s multiple-comparisons test. Statistical significance in (J) was determined by ordinary two-way ANOVA followed by Tukey’s multiple-comparisons test.

**Figure 6.**
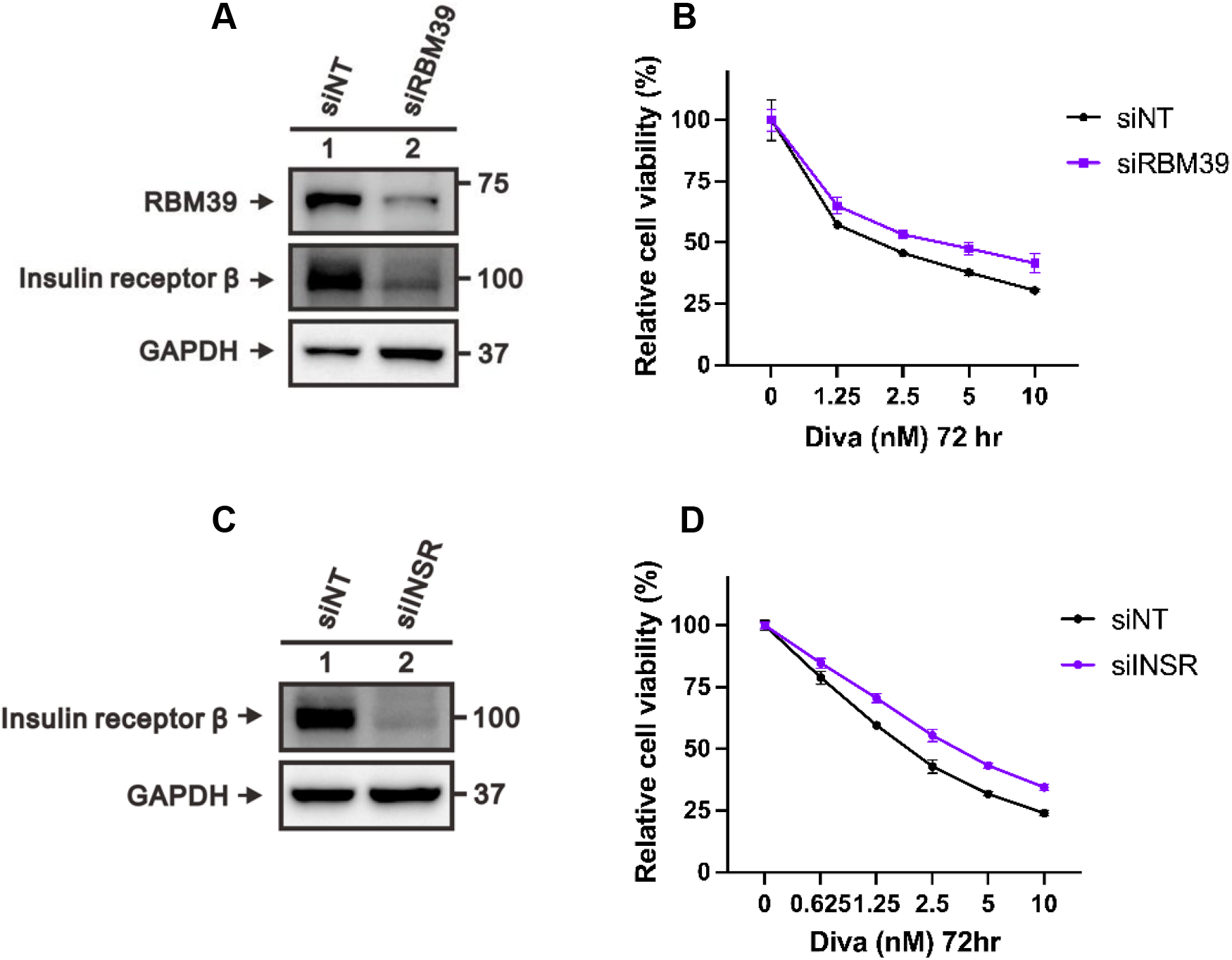
RBM39-dependent regulation of INSR contributes to divarasib-induced cytotoxicity in H358 cells. (A) Efficient knockdown of RBM39 in H358 cells transfected with RBM39-targeting siRNA (siRBM39) or non-targeting control siRNA (siNT), showing reduced RBM39 and insulin receptor β protein levels, with GAPDH as loading control, as verified by western blot analysis. (B) Cell viability of H358 cells transfected with siNT or siRBM39 and treated with increasing concentrations of divarasib (Diva) for 72 h, as determined by the CellTiter-Glo assay. (C) Efficient knockdown of INSR in H358 cells transfected with INSR-targeting siRNA (siINSR) or siNT, with GAPDH as loading control, as verified by western blot analysis. (D) Cell viability of H358 cells transfected with siNT or siINSR and treated with increasing concentrations of Diva for 72 h, as determined by the CellTiter-Glo assay. Data are presented as mean ± SD from three independent biological replicates. Statistical significance in (B) and (D) was determined by ordinary two-way ANOVA followed by Šídák’s multiple-comparisons test.

To assess INSR alternative splicing, we used a specific primer set that simultaneously detects both INSR isoforms, namely INSR-B (165 bp), which includes exon 11, and INSR-A (128 bp), which lacks exon 11 (58). We then calculated the percent spliced in (PSI) value, defined as the intensity of the inclusion band divided by the total intensity of both bands. Accordingly, a higher PSI value reflects a greater proportion of the INSR-B isoform. While divarasib treatment alone did not significantly alter the INSR isoform ratio, indisulam treatment reduced the PSI value indicating a shift toward the INSR-A isoform. Importantly, co-treatment with divarasib restored the decrease in PSI induced by indisulam (Figure 5F-H). These changes closely paralleled the alterations in RBM39 protein abundance, suggesting that RBM39 contributes to the regulation of INSR alternative splicing.

Functionally, INSR-A preferentially binds IGF-2 and promotes mitogenic signaling, whereas INSR-B primarily mediates the metabolic effects of insulin (55). To investigate the functional consequences of the altered INSR isoform ratio, H358 cells were treated with divarasib, indisulam, or their combination for 48 hours, followed by serum starvation and subsequent stimulation with either insulin or IGF-2 for 30 minutes. Compared with vehicle control, stimulation with either insulin or IGF-2 markedly increased AKT phosphorylation at Ser473, indicating successful activation of insulin receptor signaling under basal conditions (Supplementary Figure 4A–B). However, only indisulam-treated cells, which exhibited a shift toward the INSR-A isoform, showed significantly enhanced ERK1/2 phosphorylation at Thr202/Tyr204 in response to IGF-2 stimulation. Importantly, co-treatment with divarasib attenuated this increase in ERK1/2 phosphorylation (Figure 5I-J). These findings demonstrate that RBM39-dependent changes in INSR alternative splicing have functional consequences and are associated with altered ERK1/2 signaling in response to IGF-2 stimulation. Taken together, these results support a model in which divarasib-mediated stabilization of RBM39 influences both the expression and alternative splicing of the INSR gene, thereby linking RBM39 regulation to altered insulin receptor signaling and divarasib-induced cytotoxicity.

### RBM39-dependent regulation of INSR contributes to divarasib-induced cytotoxicity in NSCLC

To determine whether RBM39 itself contributes to divarasib-induced cytotoxicity, we examined the effect of RBM39 depletion on cell viability following divarasib treatment. Cells transfected with RBM39-targeting siRNA (siRBM39) exhibited a modest but statistically significant reduction in divarasib-induced cell death compared with non-targeting controls (Figure 6B), suggesting that endogenous RBM39 contributes to the downstream cytotoxic effects of divarasib. The relatively modest magnitude of this effect suggests that RBM39 is one of multiple downstream effectors involved in divarasib-induced cytotoxicity.

Since divarasib-mediated stabilization of RBM39 regulates both INSR expression and alternative splicing, we next sought to determine whether these changes contribute to the cytotoxic effects of divarasib. Consistent with our previous findings that divarasib increased INSR expression, whereas indisulam reduced it, siRNA-mediated depletion of INSR partially but significantly rescued divarasib-induced cell death (Figure 6C-D). Notably, this protective effect was consistent with that observed following RBM39 degradation by indisulam or RBM39 knockdown (Figure 3A, 3D, and 6B). These findings suggest INSR as a potential downstream effector linking RBM39 regulation to divarasib-induced cytotoxicity.

## Discussion

Our results extend the utility of stability-based proteomics beyond on-target confirmation by revealing a functional and previously unrecognized interaction between divarasib and a non-KRAS protein, RBM39. The biochemical and cellular data support a model in which divarasib stabilizes RBM39 and thereby antagonizes RBM39 degradation induced by aryl-sulfonamide molecular glues. This antagonism is recapitulated at the phenotypic level, where divarasib attenuates the cytotoxicity of three structurally distinct RBM39 degraders and, reciprocally, the degraders attenuate divarasib-induced cell death. Together with the partial protection conferred by RBM39 knockdown, these findings establish RBM39 as a functionally relevant noncanonical target of divarasib. The downstream consequences of this interaction reinforce that conclusion. Divarasib-mediated stabilization of RBM39 increased INSR expression and altered INSR exon 11 alternative splicing, with corresponding changes in insulin- and IGF-2-stimulated ERK1/2 phosphorylation. Depletion of INSR partially attenuated divarasib-induced cell death, mirroring the effect of RBM39 knockdown and identifying INSR as one downstream effector through which RBM39 influences the cellular response to divarasib. Collectively, these observations implicate RBM39 abundance, together with its regulation of the INSR signaling axis, as a potential determinant of cellular responses to both divarasib and RBM39 degraders.

This work highlights the ability of stability-based proteomics to uncover unexpected drug-protein interactions that are invisible to target-centric, adduct-based approaches. Although divarasib was designed as a highly selective covalent KRAS^G12C^ inhibitor, our findings demonstrate that it can also engage several non-KRAS proteins, with RBM39 representing a directly validated noncovalent target. This observation raises the possibility that non-KRAS engagement may represent an underappreciated feature of the broader KRAS^G12C^ inhibitor class. Future studies comparing divarasib with other clinical KRAS^G12C^ inhibitors, such as sotorasib and adagrasib, using matched stability-based proteomics and orthogonal cellular assays will help determine whether RBM39 stabilization is unique to divarasib or a shared property of covalent KRAS^G12C^ inhibitors. If so, these previously unrecognized interactions may contribute to the tumor-to-tumor variability in therapeutic efficacy and tolerability observed in the clinic.

Beyond establishing RBM39 as a divarasib-stabilized protein, our results position divarasib-mediated RBM39 stabilization as a useful chemical-epistasis tool for dissecting RBM39-degrader biology. Aryl-sulfonamide molecular glues induce cell death by recruiting RBM39 to the DCAF15 E3 ubiquitin ligase, leading to proteasomal degradation of this essential splicing factor and consequent transcriptome-wide splicing defects (28, 37, 59). Consistent with its critical role in RNA splicing, RBM39 loss promotes tumor cell death, making it an attractive therapeutic target (25). By protecting RBM39 from degrader-induced depletion, divarasib attenuates the cytotoxicity of multiple RBM39 degraders, suggesting that cellular responses to these agents are highly sensitive to RBM39 abundance. These findings are consistent with the existence of a threshold level of RBM39 below which cells undergo irreversible loss of viability, such that even modest stabilization of RBM39 can substantially blunt degrader-induced cytotoxicity. Mechanistically, our cycloheximide chase experiments indicate that divarasib counteracts indisulam-induced RBM39 degradation by increasing RBM39 protein stability. Whether this stabilization results from impaired DCAF15-dependent degradation, reduced ubiquitination, or divarasib-induced conformational changes that render RBM39 less susceptible to degradation remains to be determined. Future biochemical studies will be required to distinguish among these possibilities.

Our results showed that divarasib and indisulam exert opposite effects on INSR expression and alternative splicing. In addition, INSR knockdown partially but significantly rescued divarasib-induced cell death, identifying INSR as a potential downstream effector linking RBM39 regulation to divarasib-induced cytotoxicity. However, the molecular mechanism by which RBM39 regulates total INSR expression remains to be determined. Although we confirmed that alterations in INSR isoform composition have functional consequences on downstream signaling, whether the splicing shift itself contributes to the cytotoxicity induced by RBM39 degraders and antagonized by divarasib remains an open question. Because INSR-A preferentially mediates proliferative IGF-2 signaling, it is conceivable that degrader-induced shifts in the INSR-B/INSR-A ratio contribute to the cytotoxic program and that preservation of INSR-B by divarasib partially accounts for the rescue of cell viability. Nevertheless, the relative contributions of total INSR expression and alternative splicing to the cytotoxic effects of divarasib and RBM39 degraders remain unclear. Furthermore, RBM39 regulates a broad network of splicing events, and the modest protective effect observed following INSR depletion suggests that additional RBM39-regulated targets are likely involved. Dissecting the relative contributions of INSR expression and isoform composition, together with identifying additional RBM39-regulated targets, will require future isoform-specific and transcriptome-wide functional studies.

Our study has several limitations that define important directions for future investigation. Although two independent cysteine-reactivity datasets detected no divarasib adduct on the RBM39 cysteines they covered, three of the five RBM39 cysteines (C425, C454, and C468) were not captured in either study. Thus, covalent modification at these positions cannot be formally excluded. However, because all three residues are located well outside the RRM1 region whose folding stability is altered by divarasib, a covalent contribution to the observed RRM1 stabilization appears unlikely. Although RBM39 engagement and direct binding were demonstrated at micromolar divarasib concentrations, further studies are needed to define the concentration dependence of RBM39 engagement in cells. Additionally, our cycloheximide chase experiments demonstrate that divarasib increases RBM39 protein stability in the presence of indisulam, although the underlying mechanism remains to be determined. The contribution of altered INSR splicing to the observed phenotypes awaits INSR-isoform-specific perturbation studies. Finally, although our viability assays establish that divarasib and RBM39 degraders reciprocally modulate cell death, they do not define the underlying death program; whether these agents engage apoptotic versus other regulated cell-death pathways will require dedicated cell-death assays (60, 61). Despite these limitations and remaining open questions, protein folding stability-based proteomics identified RBM39 as a divarasib-stabilized protein, and orthogonal cellular assays demonstrated that divarasib increases RBM39 abundance, protects RBM39 from aryl-sulfonamide-induced degradation, preserves RBM39-dependent INSR expression and splicing event, and antagonizes degrader-induced cell death. Collectively, these results establish a mechanistic link between divarasib-mediated RBM39 stabilization and downstream viability responses and illustrate the power of stability-based proteomics to uncover functionally important, noncanonical targets of small molecules.

The stability-based profiling strategy applied here to divarasib is readily transferable to other KRAS-targeted agents. Indeed, we recently employed the same one-pot SPROX and TPP workflows to study ARS-1620, the prototypical KRAS^G12C^ inhibitor, and identified and validated the aldehyde dehydrogenase ALDH1A3 as a non-KRAS off-target (21). Notably, whereas the interaction between ARS-1620 and ALDH1A3 is covalent, the engagement of RBM39 reported here appears to be noncovalent. Together, these studies demonstrate that stability-based proteomic approaches can capture both covalent and noncovalent off-target non-KRAS interactions across structurally distinct KRAS inhibitors. The same one-pot SPROX workflow, together with orthogonal protein folding stability-based approaches, can also be used to identify noncovalent, non-KRAS interactors of additional clinical inhibitors, including KRAS^G12C^, KRAS^G12D^, and pan-KRAS inhibitors, and to determine whether RBM39 stabilization is unique to divarasib or represents a shared property of distinct KRAS inhibitor scaffolds. Because these compounds differ substantially in chemical structure, they are likely to exhibit both shared and compound-specific non-KRAS interactions. Comparative profiling across inhibitors, tumor types, and cellular contexts may therefore reveal previously unrecognized mechanisms of action, off-target activities, and combination liabilities. Such a comparative atlas could ultimately facilitate the rational development and clinical deployment of next-generation KRAS inhibitors.

## Methods

### One-pot SPROX

Cell pellets were resuspended in PBS solution and subjected to five freeze–thaw cycles to achieve complete lysis. The lysates were centrifuged at 14,000 × g for 15 min at 4 °C, and the resulting supernatant were collected. Protein concentrations were determined using the Bradford assay and normalized to 1.0 mg/mL. The cell lysates were then aliquoted and treated with vehicle control (DMSO), 1 µM divarasib, or 100 µM divarasib, with biological replicates as indicated in Figure 1A. Each treated lysate was distributed into PBS solutions (pH 7.4) containing increasing concentrations of urea (0.5–6 M, in 0.5 M increments). Samples were equilibrated for 4 hours at room temperature, consistent with previously reported conditions (3, 13). Following equilibration, samples were oxidized with H₂O₂ at final concentration of 1 M for 3 mins. The oxidation reaction was quenched by adding a five-fold molar excess of TCEP. Denaturant-point samples for each replicate were combined into a single “one-pot” sample, and the resulting protein mixtures were processed using the iFASP protocol as previously described (21, 24). Briefly, samples from each replicate were transferred to 10 kDa MWCO centrifugal filter unit. Buffer exchange was performed with 8 M urea in 0.1 M Tris-HCl buffer (pH 8.5), followed by reduction with TCEP (5 mM, 1 hour), alkylation with MMTs (20 mM, 5 mins), and buffer exchange into 0.1 M TEAB. Sample were then digested overnight with trypsin at a trypsin/protein ratio of 1:100. TMT 10-plex reagents were added to each sample at a 4:1 reagent/peptide mass ratio and labelling was allowed to proceed for 1 hour before quench with hydroxylamine. The resulting peptides were desalted using a C-18 column. Methionine-containing peptides were enriched using the Pi^3^ methionine reagent kit according to the manufacturer’s instructions, the samples were desalted again using a C18 column, and the resulting sample were analyzed by LC-MS/MS.

### Quantitative LC–MS/MS Analysis and SPROX data analysis

LC–MS/MS experiments were performed on a Thermo Orbitrap Exploris 480 mass spectrometer system coupled to a Thermo Easy-nLC 1200 system. Peptides were reconstituted in 1% trifluoroacetic acid (TFA) and 2% acetonitrile to 1 μg/μL, and 1 μg was injected per run. Peptides were trapped on a Thermo Acclaim PepMap 100 (75 μm × 2 cm) column and separated on a nanoViper 2Pk C18 (3 μm, 100 Å, 75 μm × 25 cm) analytical column using a 110 min linear gradient of 3–30% ACN at a flow rate of 400 nL/min and a column temperature of 45 °C. Data was acquired in data-dependent acquisition (DDA) mode with a top-20 method and 2.5 s cycle time. MS1 and MS2 scans were acquired at resolutions of 120k and 50k, respectively. The MS1 AGC target was 4.0 × 10^5^ ions with a max injection time of 10^5^ ms. HCD collision energy was set to 38%, with a scan range of 375–1500 m/z, an isolation window was 0.7 Da, and a dynamic exclusion duration of 60 s.

Raw LC-MS/MS data were searched against mammalian protein databases (UniProt Knowledgebase 2017-10-25 release) using Proteome Discoverer 2.3 (ThermoFisher) with the SEQUEST HT search engine. Search parameters included fixed MMTS modification of cysteine, TMT 10-plex labeling of lysine side chains and peptide N-termini, and variable modification for methionine oxidation, and acetylation, Met-loss and Met-loss + Acetylation of the protein N-terminus. Trypsin was specified as the protease with up to two missed cleavages allowed. Protein and peptide with an FDR confidence of “high” and “medium” (i.e., FDR < 1%) and without a single quantification channel of 0 were used for further processing.

The data analysis was similar as previous described (21). The quantified TMT intensities were normalized by the average intensities of all methionine containing peptides for each channel to correct for labeling or sample-handling variability. A Welch’s t-test was applied to the normalized data, and hits were defined as those with p-value <0.05 and an absolute log_2_ fold-change threshold of 0.3.

### Cell lines and culture

Human KRAS^G12C^-mutant NSCLC cell lines H358 and H2122 were cultured in RPMI-1640 medium (Gibco, #11875093) supplemented with 10% heat-inactivated fetal bovine serum (Sigma, #F0926) and 1× penicillin-streptomycin (Gibco, #15140122). Cells were maintained at 37 °C in a humidified incubator with 5% CO_2_. Cell line authentication was performed by short tandem repeat (STR) profiling, and all cultures were verified to be free of mycoplasma contamination prior to cryopreservation by the Cell Culture Facility.

### Cellular thermal shift assay (CETSA)

CETSA were performed as previously described (35). Briefly, H358 cells were treated with DMSO or 1 μM divarasib for 1 hour, harvested, and divided into equal aliquots. Samples were heated at temperatures ranging from 37°C to 49.5°C for 3 min and then subjected to two cycles of liquid nitrogen freeze-thaw lysis. Soluble proteins were separated from aggregated proteins by centrifugation at 16,000 rpm for 15 min at 4 °C, and the resulting supernatants were analyzed by immunoblotting for RBM39 and GAPDH, which served as a thermally stable internal control. Soluble RBM39 levels were quantified by densitometric analysis, normalized to GAPDH, and expressed relative to the 37 °C condition to generate thermal stability curves. All experiments were performed in three independent biological replicates.

### MicroScale thermophoresis (MST)

Binding of divarasib to RBM39 was quantified by MST on a Monolith instrument (NanoTemper Technologies, Munich, Germany). Recombinant N-terminally His-tagged human RBM39 (Biorbyt, cat. orb2965677) was reconstituted per the manufacturer’s instructions and exchanged into assay buffer (1× PBS pH 7.4, 0.05% Tween-20; PBS-T). Protein was labelled with the His-tag-specific RED-tris-NTA 2^nd^ generation dye (NanoTemper) and, after labelling, cleared by centrifugation to remove aggregates.

Divarasib (10 mM stock in DMSO) was serially diluted two-fold across 16 points in 100% DMSO; each dilution was then diluted into PBS-T by 100-fold to hold the solvent content constant across the series. Ligand dilutions were mixed 1:1 with labelled RBM39, giving final concentrations of 50 nM labelled target, divarasib from 50 µM to 1.53 nM, and 1% (v/v) DMSO in every sample. Samples were loaded into standard capillaries and measured at 25.0 °C using 100% excitation power and 40% MST power.

Data were analyzed in MO.Affinity Analysis v3.0.5 using the “On Time” evaluation strategy, with the cold region defined as −1 to 0 s and the hot region as 4 to 5 s. Normalized fluorescence (FNorm, ‰) was fitted to a 1:1 binding model with the target concentration fixed at 50 nM. Three independent biological replicates were performed, each with a freshly prepared dilution series; dissociation constants are reported as mean ± SD of the three independent fits.

### Western blots

Following treatment with the indicated compounds, cells were harvested and lysed in RIPA buffer supplemented with protease inhibitor cocktail (Roche, #11697498001). Lysates were incubated at 4 °C with gentle agitation for 30 min and clarified by centrifugation at 16,000 rpm for 15 min at 4 °C. Protein concentrations in the resulting supernatants were determined using the Pierce BCA Protein Assay Kit (Thermo Fisher Scientific, #23225). Samples were mixed with 4× Laemmli sample buffer (Bio-Rad, #1610747) and heated at 95 °C for 5 min before electrophoresis.

Equal amounts of protein (30–50 μg) were resolved on 10% SDS-PAGE gels and transferred onto PVDF membranes. Membranes were blocked with 5% non-fat milk in TBST for 1 hour at room temperature and subsequently incubated overnight at 4 °C with primary antibodies diluted in 5% BSA. Following three washes with TBST (10 min each), membranes were incubated with horseradish peroxidase (HRP)-conjugated anti-rabbit and anti-mouse secondary antibodies (Cell Signaling Technology, #7074 and #7076) for 1–1.5 hour at room temperature. After three additional washes with TBST (15 min each), immunoreactive signals were visualized using SuperSignal™ West Pico (Thermo Fisher Scientific, #34577) or West Femto (Thermo Fisher Scientific, #34096) chemiluminescent substrates and imaged with a ChemiDoc™ Imaging System (Bio-Rad). Band intensities were quantified by densitometric analysis, normalized to GAPDH, and expressed relative to vehicle (DMSO)-treated controls. All experiments were performed using three independent biological replicates.

Primary antibodies used in this study included RBM39 (1:1000, Invitrogen, #PA5-31103), RAS (1:1000, Cell Signaling Technology, #8955), GAPDH (1:1000, Cell Signaling Technology, #97166), β-Tubulin (1:1000, Cell Signaling Technology, # 86298), insulin receptor β (1:500, Cell Signaling Technology, #3025), phospho-ERK1/2 (Thr202/Tyr204; 1:1000, Cell Signaling Technology, #4370), ERK1/2 (1:1000, Cell Signaling Technology, #4695), phospho-Akt (Ser473; 1:1000, Cell Signaling Technology, #9271), and Akt (1:1000, Cell Signaling Technology, #9272).

### Computational docking

Glide docking was conducted using Schrödinger Maestro version 2025-4 (62), operated under M1 MacBook. The soluble NMR structure of human RBM39-U1 snRNA stem loop 3 complex (PDB 7ZAP) (36) was downloaded from the RCSB Protein Data Bank (https://www.rcsb.org/). The downloaded structure was refined using the Protein Preparation module as follows. The missing side chain was filled and unnecessary molecules including RNA were removed. H-bond assessment and restrained minimization were conducted using the default option. Divarasib was constructed using the Sketch module and prepared with the LigPrep module. OPLS4 force field was used for minimization and all possible ionized states at pH 5.4-9.4 were generated, including the original state (neutral). All generated states were used for docking. Grid was generated from the prepared protein structure using the Receptor Grid Generation module. Grid center was calculated with the adenine 104. The size of the grid box was automatically defined. The generated grid was used for SP Glide docking, performed in a default setting. 20 poses per ligand were prepared for comparison. ChimeraX-1.12 was used for visualization (63).

### Quantitative RT-PCR and analysis of INSR alternative splicing

Total RNA was isolated using the Quick-RNA Miniprep Kit (Zymo Research, #R1055) according to the manufacturer’s protocol. Complementary DNA (cDNA) was generated from purified RNA using SuperScript™ IV Reverse Transcriptase (Invitrogen, #18090200) and oligo(dT) primers. Quantitative real-time PCR (qPCR) was performed using Power SYBR™ Green PCR Master Mix (Applied Biosystems, #4368708) with gene-specific primers on a StepOnePlus™ Real-Time PCR System (Applied Biosystems). Relative transcript abundance was normalized to β-actin, and results were obtained from three independent biological replicates.

Alternative splicing of INSR exon 11 was evaluated by semi-quantitative RT-PCR using primers flanking exons 10–12, enabling simultaneous detection of the exon 11-included INSR-B isoform (165 bp) and the exon 11-skipped INSR-A isoform (128 bp) (64). PCR amplification was performed using MyTaq™ HS Red Mix (Thomas Scientific, #C755G96) and gene-specific primers. For INSR isoforms, reactions were initiated with a denaturation step at 95 °C for 10 min, followed by 40 cycles of 94 °C for 45 s, 56 °C for 45 s, and 72 °C for 20 s, with a final extension at 72°C for 10 min. β-actin was amplified under identical conditions except that 28 cycles were used with an annealing temperature of 59°C. PCR products were separated on 3% agarose gels and visualized using a ChemiDoc™ Imaging System (Bio-Rad). Band intensities were quantified by densitometry, and exon 11 percent-spliced-in (PSI) values were calculated as the intensity of the exon 11 inclusion band divided by the sum of the inclusion and exclusion band intensities. Data were derived from three independent biological replicates.

### Cell-viability assays

Cell viability was measured using the CellTiter-Glo® Luminescent Cell Viability Assay (Promega, #G7570). Cells (6–7 × 10³ per well) were seeded in white 96-well plates (Corning, #3903) and treated with divarasib, aryl-sulfonamide degraders (indisulam, E7820, or dCeMM1), or the indicated drug combinations for 72 hours. After treatment, 10 μL of CellTiter-Glo reagent was added to wells containing 100 μL of medium, and luminescence was measured after a 10-min incubation at room temperature. Cell viability was normalized to vehicle (DMSO)-treated controls. Results are representative of three independent biological experiments.

### Small interfering RNAs (siRNAs) knockdown

Gene silencing of RBM39 and INSR was achieved by siRNA-mediated knockdown. A non-targeting control siRNA (siNT) was obtained from Qiagen, whereas siRNAs targeting human RBM39 (M-011965-00-0005) and INSR (M-003014-02-0005) were purchased from Dharmacon. Reverse transfections were performed at a final siRNA concentration of 50 nM. Briefly, siRNAs were diluted in 400 μL of Opti-MEM™ I Reduced Serum Medium (Gibco, #11058021), followed by the addition of 5 μL Lipofectamine™ RNAiMAX reagent (Thermo Fisher Scientific, #13778150). After incubation at room temperature for 20 min to allow siRNA-lipid complex formation, 5-7 × 10⁵ cells were seeded directly into the transfection mixture in 6-well plates. Cells were cultured for at least 48 hours prior to downstream analyses or confirmation of knockdown efficiency.

### RNA sequencing and bioinformatic Analysis

Total RNA was extracted using the Quick-RNA Miniprep Kit (Zymo Research, #R1055), and samples (n = 3 per condition) were submitted to the Duke Sequencing and Genomic Technologies (SGT) Facility for sequencing on an Illumina NovaSeq 6000 platform. Sequencing data were analyzed by Solvuu Inc. Briefly, raw reads were trimmed using Trimmomatic (v0.39) and aligned to the human genome and transcriptome (GRCh38, GENCODE v39) with STAR (v2.7.9a). Gene-level counts, transcripts per million (TPM), and fragments per kilobase per million mapped reads (FPKM) were generated using RSEM (v1.3.3). Differential expression analysis was performed using DESeq2 (v1.28.1), and pathway enrichment analysis was conducted using Gene Set Enrichment Analysis (GSEA; v4.2.3) with the Molecular Signatures Database (MSigDB, v7.4).

Genes upregulated by divarasib treatment were intersected with a previously reported set of RBM39-regulated splicing targets (54) to identify candidate genes potentially regulated by divarasib-mediated RBM39 stabilization.

Raw and processed RNA sequencing data have been deposited in the NCBI Gene Expression Omnibus (GEO) under accession numbers GSE338666.

### The Cancer Genome Atlas (TCGA) correlation analysis

Correlation analysis between RBM39 and INSR expression was performed using the GEPIA3 web server (http://gepia3.bioinfoliu.com/), which integrates RNA sequencing data from TCGA and Genotype-Tissue Expression (GTEx) projects. Transcript abundance values were represented as log₂ (TPM + 1). Pearson correlation coefficients and corresponding p-values were calculated for each of the 33 TCGA tumor cohorts using the Gene Correlation Analysis module. Tumor types exhibiting significant positive correlations were summarized, and representative scatter plots for lung adenocarcinoma (LUAD) and lung squamous cell carcinoma (LUSC) were generated using the same platform.

### Statistical analysis

Data are presented as mean ± standard deviation (SD) from three independent biological replicates. Individual biological replicates are shown in bar graphs, whereas the number of biological replicates for line graphs is specified in the corresponding figure legends. Statistical analyses were performed using GraphPad Prism software. Differences among groups were assessed using ordinary one-way analysis of variance (ANOVA) followed by Tukey’s or Dunnett’s multiple-comparisons test, or ordinary two-way ANOVA followed by Šídák’s or Tukey’s multiple-comparisons test, as appropriate. Error bars represent SD, and exact p-values are provided in the figures when applicable. Additional statistical details, including sample size and significance thresholds, are described in the corresponding figure legends.

## Supporting information

Supplementary reagent list

## Acknowledgments

This work was funded in part by a National Institutes of Health grant (1R21-CA297690-01A1) to J-T.C. and M.C.F.

## Author Contributions

S-Y.C., Y.Z., M.C.F., and J-T.C. conceived the experiments and wrote the manuscript. Initial experiments were performed by J.W. S-Y.C. and Y.Z. performed the majority of the experiments. J.H.S., J.H., M.C.F., and J-T.C. supervised the work. J.W., G.N., H.L., Y.C., C.F., Y.S., C-C.L., and S-C.W. collaborated in the discussion and experiments.

## Data and Code Availability

The mass-spectrometry proteomics data have been deposited to the ProteomeXchange Consortium via the PRIDE partner repository (65) with the project accession (PXD079782) and token (zuzglHB7f8tN); RNA-seq data have been deposited in the NCBI Gene Expression Omnibus (GEO) under accession numbers GSE338666. Any analysis code will be listed in the Key Resources Table and made available before publication.

