## Supplementary figures for "The KRAS^G12C^ Inhibitor Divarasib Stabilizes RBM39 and Antagonizes Aryl-Sulfonamide Degraders"

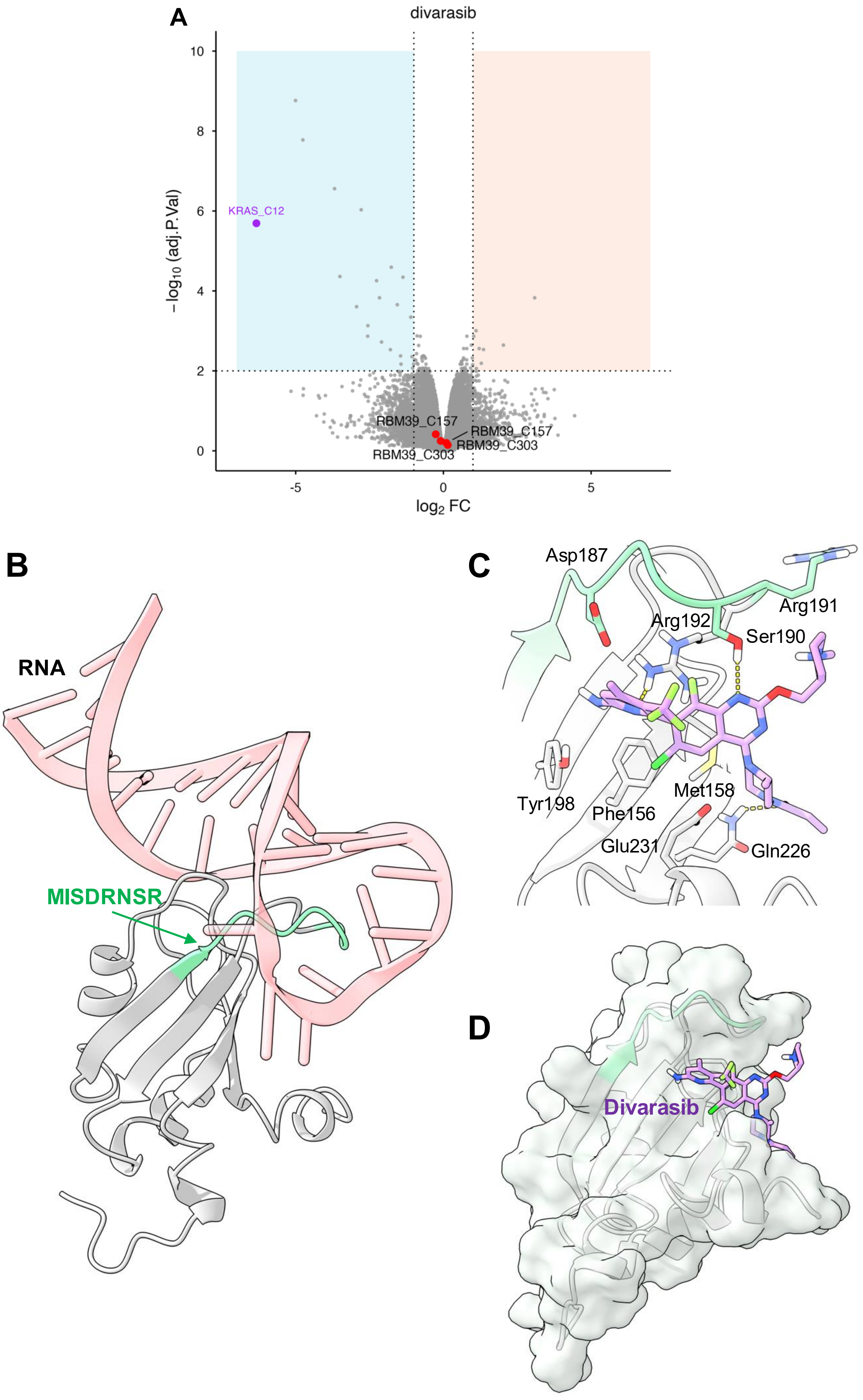


**Supplementary Figure 1. Computational analyses of divarasib engagement with RBM39**

(A) Reanalysis of divarasib cysteine-reactivity profiling data reported by Condakes et al. (34). Dots in the blue quadrant represent cysteine-containing peptide fragments found to be significantly reduced in samples treated with inhibitor ($\log_2 FC < -1$ and adj. $p$-value $< 0.01$); dots in the red quadrant represent cysteine-containing peptide fragments found to be significantly exposed in samples treated with inhibitor ($\log_2 FC > 1$ and adj. $p$-value $< 0.01$). The fragment for $KRAS^{G12C}$ is highlighted in purple. The fragments for RBM39 are highlighted in red. (B-D) Docking model of divarasib with RRM1 domain of RBM39. (B) NMR spectroscopy structures of RRM1 (PDB ID: 7ZAP), shown with the bound RNA, which was removed prior to docking. The SPROX-responsive peptide is highlighted in green. (C) Docking model of divarasib. Divarasib is represented as purple stick. Hydrogen bonds are represented in yellow dashes. (D) Surface representation of RRM1 domain of RBM39 (gray).

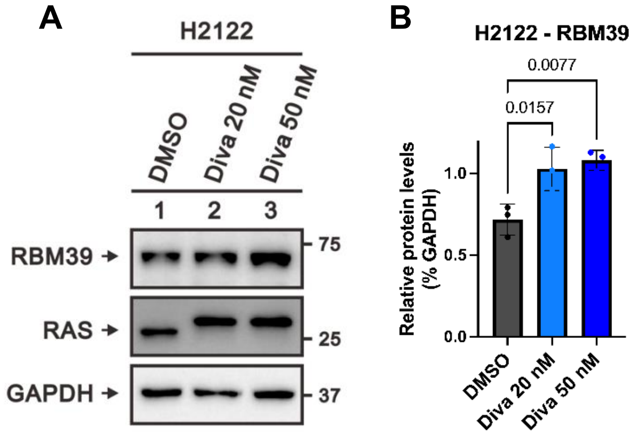


**Supplementary Figure 2. Divarasib increases basal RBM39 protein abundance in H2122 cells**

(A, B) RBM39 protein levels in H2122 cells treated with DMSO or divarasib (Diva; 20 or 50 nM) for 72 h. (A) Representative immunoblots of RBM39, RAS, and GAPDH. (B) Quantification of RBM39 protein levels normalized to GAPDH. Data are presented as mean ± SD from three independent biological replicates. Statistical significance in (B) was determined by ordinary one-way ANOVA followed by Dunnett's multiple comparisons test.

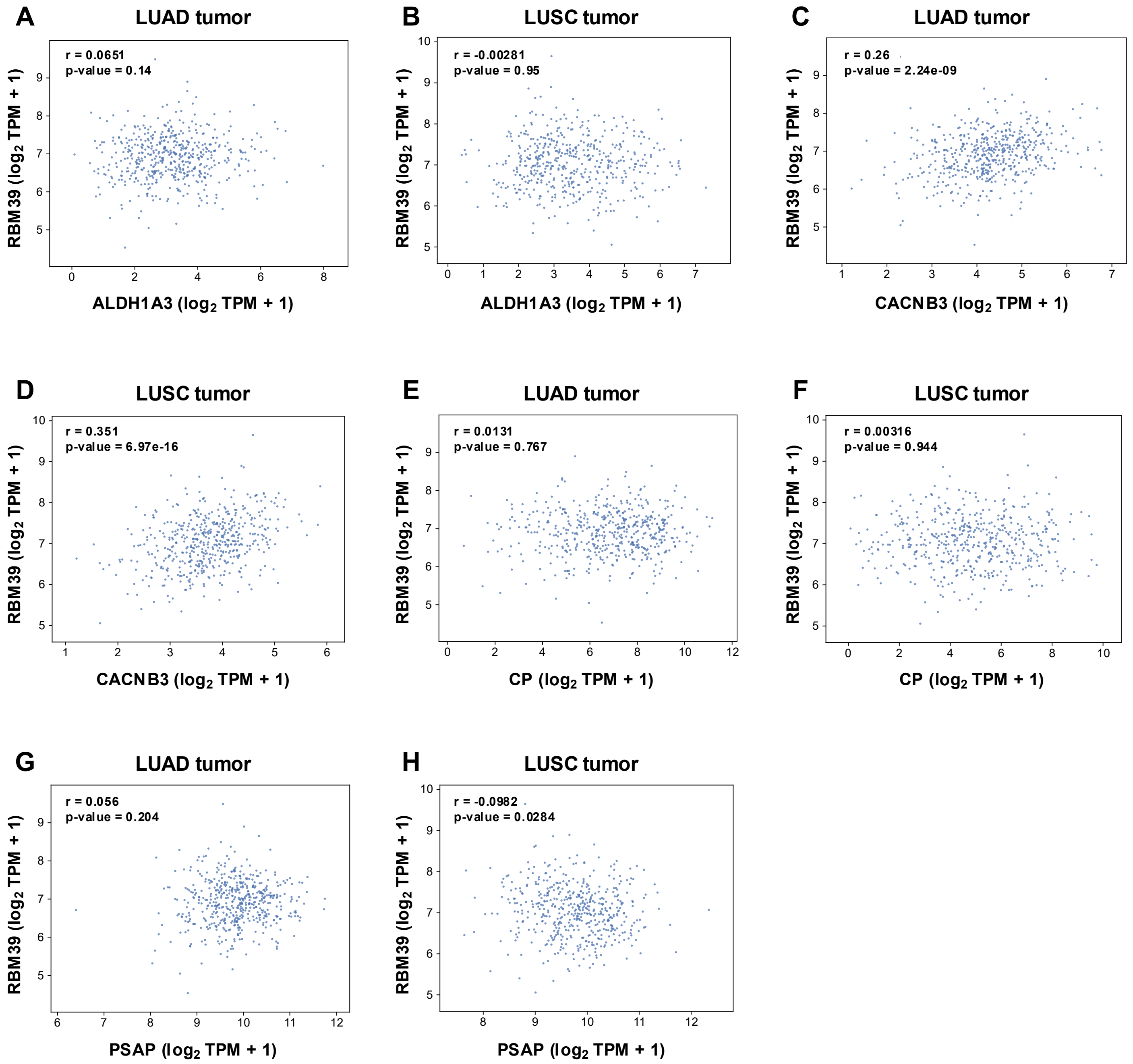


**Supplementary Figure 3. Correlation analysis of overlapped genes with RBM39 in LUAD and LUSC from TCGA datasets**

(A, B) Scatter plots showing the correlation between RBM39 and ALDH1A3 transcript levels (log2 TPM + 1) in (A) lung adenocarcinoma (LUAD) and (B) lung squamous cell carcinoma (LUSC). (C, D) Scatter plots showing the correlation between RBM39 and CACNB3 transcript levels (log2 TPM + 1) in (C) LUAD and (D) LUSC. (E, F) Scatter plots showing the correlation between RBM39 and CP transcript levels (log2 TPM + 1) in (E) LUAD and (F) LUSC. (G, H) Scatter plots showing the correlation between RBM39 and PSAP transcript levels (log2 TPM + 1) in (G) LUAD and (H) LUSC. Data shown in (A–H) are based on data generated by the TCGA Research Network (https://www.cancer.gov/tcga).

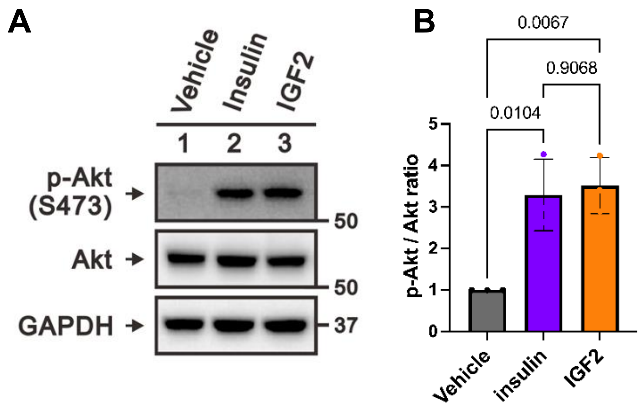


**Supplementary Figure 4. Insulin and IGF-2 stimulate Akt phosphorylation in H358 cells.**

(A, B) Akt phosphorylation at Ser473 in H358 cells following 6 h of serum starvation and subsequent stimulation with insulin (100 nM) or IGF-2 (100 ng/mL) for 30 min. (A) Representative immunoblots of phospho-Akt (Ser473), total Akt, and GAPDH. (B) Quantification of phospho-Akt (Ser473) levels normalized to total Akt. Data are presented as mean ± SD from three independent biological replicates. Statistical significance in (B) was determined by ordinary one-way ANOVA followed by Tukey's multiple-comparisons test.
